# The involvement of type IV pili and phytochrome in gliding motility, lateral motility and phototaxis of the cyanobacterium *Phormidium lacuna*

**DOI:** 10.1101/2021.03.22.436410

**Authors:** Tilman Lamparter, Jennifer Babian, Katrin Fröhlich, Marion Mielke, Nora Weber, Nadja Wunsch, Finn Zais, Kevin Schulz, Vera Aschmann, Nina Spohrer, Norbert Krauß

**Author notes:** These authors contributed equally to this work (Jennifer Babian, Katrin Fröhlich, Marion Mielke, Nora Weber, Nadja Wunsch).

## Abstract

*Phormidium lacuna*, a filamentous cyanobacterium without heterocysts, can be transformed by natural transformation. These filaments are motile on agar and other surfaces and display rapid lateral movements in liquid culture. Furthermore, they exhibit phototactic response under vertical illumination in Petri dishes. We generated mutants in which a KanR resistance cassette was integrated in the phytochrome gene *cphA* and in various genes of the type IV pilin apparatus. *pilM, pilN, pilQ* and *pilT* mutants were defective in all three responses, indicating that type IV pili are involved in all three kinds of motility. Rapid movements of wild type in liquid culture requires an extracellular matrix with type IV pili as central player. *pilB* mutants are only partially blocked in their responses. *pilB* is the proposed ATPase for expelling of the filament. In the mutant, this function could be overtaken by an alternative protein, like *pilT*, which regularly mediates retraction of pili. The *cphA* mutant revealed a significantly reduced phototactic response towards red light. We assume that together with other photoreceptors, CphA regulates the phototactic response by down regulation of surface attachment.

## Introduction

Around 3.0 billion years ago, cyanobacteria evolved oxygenic photosynthesis [1]. This invention increased biological growth drastically caused a rise in atmospheric oxygen that changed the diversity and complexity of species on earth dramatically. All photosynthetic eukaryotic organisms are descendants of a eukaryote with a cyanobacterial endosymbiont. [2, 3]. All life on earth depends on oxygenic photosynthesis that arose in cyanobacteria. Four main kinds of cellular organizations are realized in cyanobacteria, single celled species, linear filaments, linear filaments with heterocysts or branched filaments.

Single celled and filamentous cyanobacteria can move by a gliding mechanism on surfaces [4–6]. In members of the Nostocales [7], i.e. filamentous cyanobacteria with heterocysts, this motility is often restricted to hormogonia [4, 8], a cell type formed during environmental stress. Most genetic and molecular studies on cyanobacterial motility have been performed with single celled species *Synechocystis* PCC6803 and *Synechococcus elongatus* PCC 7924 [9–11] and one member of Nostocales [7] *Nostoc punctiforme* [8]. The *Synechocystis* PCC 6803 studies involvement of type IV pili in gliding movement [9]. Type IV pili as driving complex for gliding (often: twitching) motility have intensely been investigated in other bacteria such as *Pseudomonas aeruginosa* [12] or *Neisseria meningitis* [13]. The type IV driven movement is based on a retraction and extrusion of protein filaments. The PilA protein, formed from the PilA precursor protein *via* cleavage by the protease PilD, concatenates to build extracellular pili that are expelled or retracted by motive force generated inside the cell. The ATPases PilB and PilT function as motorproteins for expelling and retraction, respectively, and PilQ forms channels for translocation of PilA through the outer membrane [14, 15]. PilM and PilN are involved in the assembly process at the inner or outer side of the inner membrane, respectively [16]. Depending on the organism, there are also several other Pil proteins that contribute to channel formation or assembly [16].

Cyanobacteria can have several PilA homologs. In *Synechocystis* PCC 6803, PilA1 is required for motility as gene knockouts result in a nonmotile phenotype. There are 9 PilA homologs in *Synechocystis* PCC 6803 with either indirect roles in motility or unknown functions [17, 18]. Knockouts of *pilC, pilD* and *pilT1* also resulted in a nonmotile phenotype, whereas knockouts of *pilA2* and *pilT2* were still motile [9]. For *Nostoc punctiforme* it was found that motility is lost in mutants in which *pilB, pilN* or *pilQ* were defective [8]. Type IV pili are also regarded as a machinery for DNA uptake in natural transformation [16, 19, 20] and in this context, cyanobacterial genomes have recently been screened for pilin genes [19, 21, 22]. Almost all out of 400 analyzed cyanobacteria have a complete set of type IV pilin genes. This wide distribution suggests that type IV pili could be the apparatus for gliding motility in all cyanobacteria, although in filamentous species belonging to the order Oscillatoriales [7] have as yet not been analyzed. However, other mechanisms have been considered for movement of Oscillatoriales [23]. Although Oscillatoriales are a major cyanobacterial group, only few genetic and molecular studies have been performed. For these reasons we considered that more studies on the functions of type IV pili should be performed.

What is the evolutionary background of the gliding movement of cyanobacteria? Many cyanobacteria move either towards the light or away from the light in a process termed positive or negative phototaxis. This effect maximizes photosynthetic light capture or protects the cyanobacterium from damage of the photosynthetic apparatus if the light is too strong, respectively. The molecular background of cyanobacterial phototaxis has again been most thoroughly investigated for the model species *Synechocystis* PCC 6803 [24–26]. Phototaxis requires a photoreceptor and a mechanism to transform the light direction into an intracellular gradient of signal transmitting molecules. In *Synechocystis* PCC6803, a light focusing effect results in a high light intensity on the light avoiding side of the cell [26]. The photoreceptor is probably located at the periphery of the cell. This focusing mechanism has been confirmed for *Synechococcus elongatus* PCC 7924 [11]. In *Synechocystis* PCC 6803, several chromoproteins such as Cph2, PixJ and PixE can serve as photoreceptors for phototaxis. PixE is a flavin-binding BLUF protein, Cph2 is a phytochrome-like protein and PixJ a cyanobacteriochrome. Regular cyanobacterial phytochromes such as Cph1 of *Synechocystis* PCC 6803 [27], have not been reported to be involved in phototaxis. In cyanobacteriochromes, phytochrome-like proteins and phytochromes, a bilin chromophore is bound to a GAF domain [28]. This chromophore undergoes light-triggered changes between two stable spectral forms. In phytochromes these are red- and far-red absorbing, whereas in cyanobacteriochromes, different spectral forms are realized. The groups differ also in their domain organizations. A regular phytochrome consists of PAS, GAF and PHY domains and a C-terminal output module such as a histidine kinase. Cyanobacteriochromes lack the PAS or the PHY domain or both, but carry other domains in high variability. Interestingly, there are often several GAF domains united in one protein, such as in Cph2. A phototaxis photoreceptor of *Synechococcus elongatus* is a PixJ homolog which contains 5 GAF domains and which is located at the periphery of the cell [11].

Our group has isolated *Phormidium lacuna* from marine rockpools, characterized growth and other parameters and sequenced the genome. *Phormidium lacuna* is a filamentous species without heterocysts that belongs to Oscillatoriales. According to 16S rRNA based phylogenetic studies [22], this species is located in the clade C4 according to [29]. *Phormidium lacuna* is quite distantly related to the model species *Synechocystis* PCC6803 or *Nostoc punctiforme*. Directly after isolation, it became obvious that *Phormidium lacuna* filaments are motile on agar surfaces, but a phototaxis response was not observed. *Phormidium lacuna* contains one typical phytochrome with PAS, GAF and PHY domains and 16 proteins that have one or more GAF domains with chromophore binding cysteine residue [30]. Five of these reveal close partial BLAST homology with PixJ of *Synechocystis* PCC6803 or *Thermosynechococcus elongatus. Phormidium lacuna* harbors also all genes for type IV pili [22].

Molecular studies on filamentous cyanobacteria are often hampered by the difficulty to perform genetic manipulations. During trials for gene transformation we found that *Phormidium lacuna* cells can take up DNA naturally and integrate it into chromosomes by homologous recombination. This was the first successful trial for natural transformation of an Oscillatoriales member [22]. After this discovery we aimed at disruption of pilin genes and the phytochrome gene in *Phormidium lacuna* by insertional mutagenesis. We made disruptant mutants of *pilA1, pilB, pilD, pilM, pilN, pilQ* and *pilT*, and the phytochrome gene *cphA*. These lines were investigated for longitudinal movement on agar and lateral movement in liquid medium. During our studies we found experimental conditions for phototaxis, so that we could compare the phototactic response of mutants and wild type. Type IV pili are clearly involved in all three kinds of movements of *Phormidium lacuna* and CphA seems to play a photoreceptor role in phototaxis and surface attachment.

## Methods

### Culture of *Phormidium lacuna*

The strain HEDO10 of *Phormidium* lacuna was used for all experiments. The genome of the strain HE10JO10 has been sequenced earlier. The recently sequenced genome of HE10DO differed in only ca. 1000 bases from the published HE10JO sequence. The filaments were cultivated in f/2 seawater medium [30, 31] in 20 ml or 50 ml culture flaks under continuous white LED light of 300 μmol m^−2^ s^−1^ under agitation at 25 °C.

### Construction of disruption vectors and transformation

The construction of the *chwA* mutant (the gene was previously termed 7_37) was described earlier [22]. For construction of the other mutants, ca. 2000 bp of each coding sequence (or when necessary, upstream sequence) were amplified from genomic DNA by PCR using the primers as listed in Table S1. Each sequence was cloned into the pGEMT easy *E.coli* vector (Promega, Madison, WI, USA) and the plasmid linearized by PCR using primers that bind to the center of the cloned sequence (Table S1). The KanR cassette was cloned into this sequence by sticky end cloning using restriction enzymes that can be recognized in the primer sequences. The positions where the sequences are interrupted 3’ of the start codon are given in Table 1. For transformation of *Phormidium lacuna*, the respective *E.coli* vector was purified by midi prep (Macherey Nagel, Düren, Germany). *Phormidium lacuna* was cultivated in 100 ml f/2 medium until A_750 nm_ reached 0.35 (ca. 5 d). The cell culture was centrifuged at 6000 × g for 3 min and the supernatant was discarded, with a small portion of the supernatant the cell volume was brought to 800 μl. Ten μg vector DNA were mixed with 100 μl cell suspension and pipetted in the center of an f/2 agar plate (1.5 % Bacto agar and 75 μg/ml Kn). This procedure was repeated to get 8 Petri dishes with DNA and *Phormidium cells*. The Petri dishes were closed with parafilm. Subsequent steps were dependent on the filament growth. Typically, the filaments were transferred after 2 weeks to plates with 500 μg/ml kanamycin (Kn) and kept for another 2 weeks on Kn medium. Thereafter, single growing filaments were isolated and cultivated on fresh agar plates with 500 μg/ml Kn. Propagation continued in liquid medium with 100 μg/ml Kn. To test for integration into the homologous site and completeness of segregation, we used inner and outer primer pairs. The inner primer pair binds at the 5’ and 3’ ends of the integrated sequence, the outer primer pair binds just 5’ and 3’ outside the integrated sequence, respectively (see Table S1 for primers). When segregation was complete (no more wild type band visible), the mutant was used for physiological assays. Segregation of *pilA1’* was incomplete over the entire observation time of several months and during the motility experiments. Disruption of *pilD* was not possible. Details are given in the Results section. For cultivation, mutants were always grown in medium with 100 μg/ml Kn, for experiments the cells were always transferred into medium without antibiotics (Supplementary Fig. S1 for PCR data).

**Table 1:** *Phormidium lacuna* proteins and genes addressed in the present study. In the third column, the lengths of the protein sequences and the identities with *Synechocystis* PCC 6803 protein homologs [50] are given. The fourth column gives the (proposed) position of the insertion of the KanR resistance cassette and the length of the entire open reading frame (ORF) in the number of basepairs.

| Protein name | Proposed function | # amino acids/<br>identity with<br>*Synechocystis* | Position of<br>integration /<br>length of<br>ORF in # bp |
| --- | --- | --- | --- |
| ChwA | Hypothetical, ChW repeats |  | 531 / 1143 |
| PilA1 | Precursor for pili protein | 173 / 39 % | 19 / 513 |
| PilB | ATP ase motor protein,<br>extension | 676 / 63 % | 36 / 2016 |
| PilD | PilA peptidase, inner membrane | 271 / 50 % | 89 / 252 |
| PilM | Pilin pore forming, inner<br>membrane | 373 / 55 % | 27 / 1116 |
| PilN | Pilin pore stabilizing inter membrane | 306 / 23 % | 480 / 783 |
| PilQ | Pore forming outer membrane | 909 / 35 % | 1081 / 2532 |
| PilT | ATP ase motor protein, retraction | 379 / 69 % | 21 / 948 |
| CphA | Photoreceptor | 764 / 44 % | 1103 / 2292 |

### Motility experiments

For all experiments on motility, *Phormidium lacuna* was transferred to Kn-free liquid f/2^+^ medium and cultivated for 5 days until A_750 nm_ reached 0.35 −0.4. The PCR pattern of the integration site (Supplementary Fig. S1) remained unchanged during this non-selective growth. The filaments were then treated with an Ultraturrax rotating knife (Silent Crusher, Heidolph, Schwabach, Germany) at 10000 rpm for 3 min. For motility studies on agar, filaments were usually concentrated 10-fold by centrifugation and 100 μl of the solution were dispersed on a Bacto agar f/2 Petri dish of 5 cm diameter. Images were recorded in 1 min intervals using a conventional microscope and a Bressler (Rhede, Germany) ocular “full HD” camera. The intensity of the microscope light was 250 μmol m^−2^ s^−1^. In some experiments, the Ultraturrax treated filaments were directly pipetted onto a 5 cm petri dish with agar medium, kept for 4 h in a growth chamber and photographed through a microscope.

For studying movement in liquid f/2 culture, 8 ml filament suspension (OD _750 nm_ = 0.4) were transferred in a 5 cm Petri dish without agar. The movement of the filaments was observed through a Leica DM750 microscope and recorded with an EC3 camera for 30 s (no time lapse). Of each sample, recordings at 3 to 10 different characteristic positions were taken.

For phototaxis experiments, we constructed a plastic holder in which a 5 mm LED is mounted at one end of a 15 mm long vertical tube with the light beam pointing upward. For blue, green and red light, LEDs with maximum emissions at 470 nm, 520 nm and 680 nm were chosen, respectively. The light intensity was 15 μmol m^−2^ s^−1^ unless indicated otherwise. Eight ml Ultraturrax treated *Phormidium* suspension (OD _750 nm_ = 0.4) were pipetted into a 5 cm Petri dish which was then placed on the LED-holder so that the LED is in the middle of the petri dish. The samples were kept for 1, 2 or 3 d in a dark room. Typically, the LED irradiation results in the formation of a circle of filaments in the center of the Petri dish. After LED irradiation, the Petri dish was photographed at its original position, a movement of the Petri dish could partially destroy the circle. The Petri dish was then shaken manually for 3 s and photographed again. The diameters of the circles were measured with ImageJ (NCBI). To this end, the image was loaded with ImageJ and a measuring line was drawn through the center of the petri dish. From the profile of the line, the diameter of the central circle was determined by the distance between halfmaximal changes of pixel intensities. Sometimes filaments formed aggregates that were not the result of phototaxis. These were not considered in our evaluations. If only aggregates were observed and no evidence for phototaxis was found, the diameter of sample was recorded as zero. Phototaxis based aggregation could be clearly distinguished from other aggregations by the position within the Petri dish and by the fact that only phototaxis based aggregation results in surface attachment.

## Results

### Generation of insertion mutants

Type IV pili are involved in gliding motility of many bacteria including single celled cyanobacteria and members of the Nostocales (filamentous species with heterocysts). To see how type IV pili are involved in a member of Oscillatoriales (filamentous species without heterocysts), we aimed at disrupting genes of PilA1, PilB, PilD, PilM, PilN, PilQ and PilT in the genome of *Phormidium lacuna*. The sequences were identified as BLAST homologs of *Synechocystis* PCC 6803 and other bacteria [22]. The degree of homology between *Synechocystis* PCC 6803 and *Phormidium lacuna* proteins is given in Table 1. We also disrupted the phytochrome gene of *Phormidium lacuna* in order to test for a possible role of this photoceptor in phototaxis. Phytochromes are abundant in most cyanobacteria, but their biological role in these organisms is unclear [32]. *Phormidium lacuna* contains one phytochrome, which is denominated CphA here, in accordance with CphA from *Fremyella diplosiphon* [33]. In the present study, we did not address other possible photoreceptor proteins such as Cph2 [10], PixJ [11, 34] or BLUF proteins [10].

As a control for possible effects on the KanR cassette we used a mutant in which the gene of ChwA is disrupted. *chwA*, originally termed 7_37 (open reading frame 37 on scaffold 7), was the first gene of *Phormidium lacuna* for which natural transformation and homologous integration of a KanR cassette was successful [22]. The protein is annotated in the NCBI database as hypothetical protein. It has 3 ChW [35] repeats at its C-terminus, hence the name ChwA in the present study. A significant effect of the KanR resistance cassette was considered because the *chwA* mutant grows at Km concentrations of more than 20 mg/ml. This very strong resistance was also found for all other lines in which KanR was integrated into the genome. Although physiological experiments were performed on Kan-free medium, expression of KanR could have an impact on motility.

For gene disruptions, cloning vectors were constructed that carried 2000 bases of the gene of interest or upstream of it. The KanR cassette was placed in the center of this sequence, so that there are 1000 bp of homologous sequence on both sides of the resistance cassette.

In this way, we built disruption vectors for *pilA1, pilB, pilD, pilM, pilN, pilQ, pilT* and *cphA* and obtained resistant transformants for all insertion constructs. For *pilB, pilM, pilN, pilQ, pilT* and *cphA* genes, the integration into the expected site and complete segregation into all chromosomes was confirmed by PCR using inner and outer primers that bind to chromosomal sequences inside and outside the integration vector, respectively (Supplemental Table S1 for list of primers and Supplemental Figure S1 for PCR with outer primers). The PCR pattern remained during 7 days on non-selective medium, the maximum time that was used for physiological experiments. In case of *pilA1*, 7 out of 8 resistant lines showed a PCR double band indicative of partial integration, *i.e.* one part of chromosomes with interrupted sequences and the other part with wild type pilA1 (Supplemental Figure S1). The incomplete segregation pattern remained unchanged over many months in selective growth medium. Complete segregation of pilA1 insertion is thus impossible, indicating that the complete loss of pilA1 would be lethal. Although only limited information can be gained from such mutants, we performed also experiments with one of these strains. This line is termed *pilA1’*. Another resistant line had only the wild type *pilA1* band. In this line, the KanR sequence must have integrated into another site. All these mutants show that disruption of pilA1 and complete segregation is lethal.

*pilD* knockout trials yielded only two resistant lines out of 4 transformations, although the same transformation effort was undertaken as with the other genes. During the first two weeks of selection, both mutants grew very slowly, but after another 2 weeks, the filaments recovered and grew at normal speed. In these lines, the pilD gene was not interrupted. We assume that the slow growth is caused by disruption of *pilD* genes in a certain proportion of the chromosomes and that by a second recombination event, the KanR cassette moved to another site. This suggests that the loss of *pilD* is also lethal. The *pilD* knockout of *Synechocystis* PCC 6803 is also lethal, but the cells can survive on glucose [36]. We did not try selection on glucose medium.

### Motility on agar

The motility on agar of *Phormidium* was usually investigated with filaments that were treated with an Ultraturrax rotating knife. This treatment separates adjacent filaments of the liquid preculture and cuts long filaments into shorter ones. Filaments were then brought onto agar surface and images were recorded in 1 min intervals through a microscope for several h.

After placement on the agar surface, all wild type filaments started to move instantaneously. The direction of movement was always longitudinal towards one end of the filament. The polarity of movement, indicating whether a filament moves towards one or the other tip, switched frequently in an apparently random manner (movie 1). However, the movement of the filaments was not completely straight. The filaments were often bent and each filament could change the direction of its movement within the laboratory coordinate system continuously. Each filament left a visible trace of extruded material on the agar surface which could serve as guide for other filaments or the same filament when it returned to the same position in a circle. When a filament touched another filament, it aligned its longitudinal direction so that finally long parts of both filaments were parallel and adjacent. Generally, filaments remained together, and additional filaments joined so that bundles of many filaments were formed (movie 1 and Fig. 1A-E). These filament bundles were usually one cell layer thick, sometimes a second layer of few filaments located above the lower one. More than two layers were not observed. The parallel leaf like filament arrangement of the bundles was highly flexible. All filaments moved continuously, also versus each other, bundles joined and separated, and higher ordered structures like circles were formed. Clearly, bundle formation and bundle dynamics are important functions of filament movement and communication between the filaments is required for such coordinated movements. Most of these characteristics can be observed in the time lapse movie 1 and in Fig. 1.

**Figure 1.**
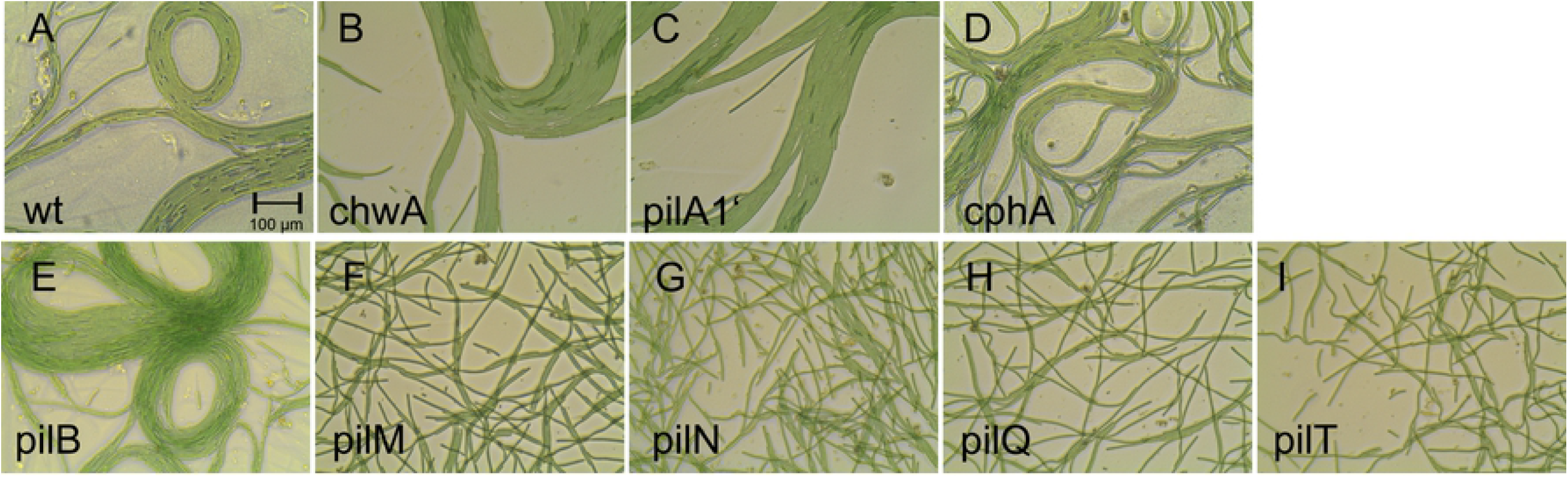
*Phormidium lacuna* filaments were treated with an Ultraturrax and placed for 4 h on agar surface. The name of each mutant is given in each panel.

The speed of movement of wild type and mutant filaments was estimated from data obtained during the early stage after Ultraturrax treatment, during which single filaments or few parallel filaments predominate. The distances covered during 1 min were analyzed for 50 to 150 filaments of each line. During this short time interval, the switching of direction is rare and the measurements are thus more reliable than for longer time intervals. As can be seen in the histogram in Fig. 2 A, the speed of wild type filaments was variable between 2 and 60 μm / min. The interval between 0 and 2 μm could not be resolved under these conditions. About 30 % of wild type filaments were in the range from 5 to 10 μm, the range that contained most filaments. The mutants could be divided into two groups, one group consisting of *chwA, pilA1’* and *cphA* which showed slightly reduced wild type like movements (Fig. 2 B-E) and the other group comprising *pilB, pilM, pilN* and *pilQ* which showed drastically reduced movements or no movements. Since none of the data follow a Gaussian distribution, we estimated the significance of differences by the non parametric Kruskal Wallis ANOVA test [37]. In this way, results from all mutants were compared with each other. All cell lines were found to be significantly different from each other (p < 0.05) and from the wild type with the exceptions of the three pairs *chwA-cphA, chwA-pilA1’* and *cphA-pilA1’*, for which the differences were not significant. This is indicated by the lines between panels in Fig. 2 B,C,D,G,H. As noted above, the speeds of *chwA, chpA* and *pilA1’* were reduced as compared to the wild type. The most likely reason for this common feature is an effect of the KanR cassette on motility. The resistance cassette seems to slightly reduce the motility. Note that during these tests all filaments grew on Kn free medium.

**Figure 2.**
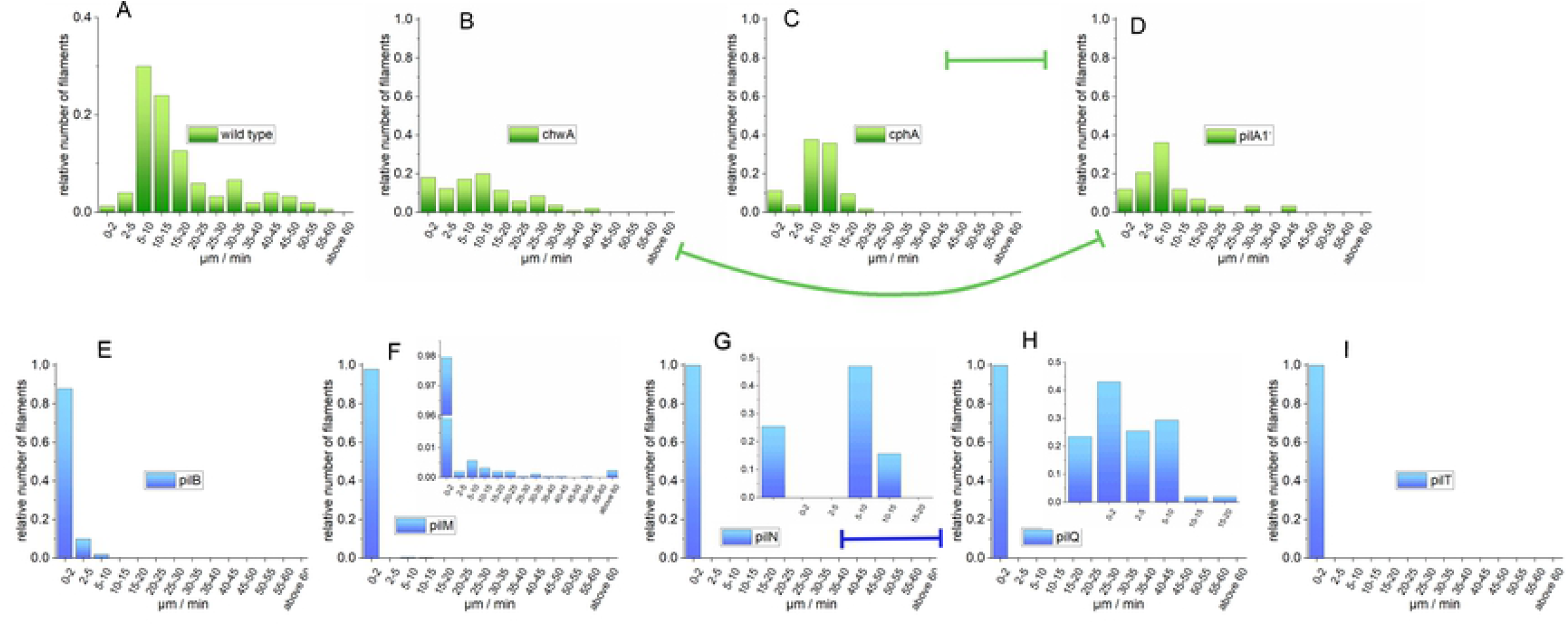
Histograms of motilities of wild type and mutant filaments after Utraturrax treatment on agar surface. Results of wild type (A), *chwA* (B), *pilA1’* (C) and *pilA1’* are represented by green bars. These lines showed a clear motility on agar. Results of *pilB* (E), *pilM* (F), *pilN* (G), *pilQ* (H) and *pilT* (I) are represented by blue bars. In these lines motility is drastically restricted or zero. The inset in F shows the results of *pilM* with expanded Y-axis. The insets in G and H for *pilN* and *pilQ* are based on 1 h measurements that were calculated back to 1 min. The 1 h measurement of *pilT* did not indicate any movement and is not shown. The lines with blocked ends are drawn between pairs of mutants that have no significantly different data (B-D, C-D, G-H), those of all other pairs of mutants or wild type are significantly different (Kruskal Wallis test, p < 0.05).

The mutants *pilB, pilM, pilN, pilQ* and *pilT* exhibited significantly reduced motility. This shows clearly that gliding motility is based on the action of type IV pili as shown for cyanobacteria of section I [9] and section IV [8], and for many bacteria [12]. The pilus motor proteins PilB and PilT, the inner membrane pore forming proteins PilM and PilN and the outer membrane core protein PilQ are all required for proper functioning of type IV pili and for twitiching motility of *Phormidium lacuna*. However, only *pilT* mutants have completely lost their motility, whereas in all other mutants there was a small fraction of filaments that remained motile or at least slightly motile. In case of *pilB* (Fig. 2 E), the fraction of motile filaments was ca 10 %. Among the pilin mutants, this was the one with the highest motility. For *pilM*, a fraction of 4% remained motile with a distribution like the wild type. This is shown in the inset of Fig. 2 F. For *pilN* and *pilQ*, the movement was also measured during 1 h time intervals. In this way also slow movements were detected (inset of Fig. 2 G and H). Such 1 h measurements were performed with *pilT* as well, but no movement was detected at all.

Figure 1 shows Ultraturrax treated filaments that were kept for 4 h on agar. Wild type filaments aligned parallel to each other as described above, forming bundles of up to 20 (Fig. 1). Bundles were also formed by *chwA, pilA1’* and *cphA*, and no difference to the wild type was observed. With *pilB*, also bundles were formed that could not be distinguished from the wild type. As noted above, only a fraction of 10% of *pilB* filaments seems to move (Fig. 2), but the long term observations ss in Fig. 1 show that all filaments of pilB can move. We confirmed a “wake up” of previously immobile *phrB* filaments when time lapse recordings were re-investigated. *pilM, pilN, pilQ* and *pilT* did not form bundles but filaments remained single or crossed over. Since in the pilin mutants there is a clear correlation between loss of bundle formation and reduced or absent motility, the gliding motility based on the type IV pili must be the motor for bundle formation.

### Motility in liquid medium

A largely unnoticed kind of movement that is different from the gliding movement has been described for filamentous cyanobacteria [38]. This movement occurs in liquid culture in lateral direction, *i.e.* perpendicular to the filament axis. We observed such movements when *Phormidium lacuna* samples were transferred from liquid culture into Petri dishes for observation with the microscope. The lateral movement was more rapid than the longitudinal gliding movement described above. It could be observed directly without time lapse recordings. When Ultraturrax treated filaments were observed in liquid culture, some filaments started immediately with lateral movements, and within few minutes, hedgehog-like bushes were formed, with ends of the filaments pointing away from the center towards the medium. Also, strands of several filaments were formed, connecting bushes with each other. Besides the winking-like, lateral movement of single filaments, groups of filaments (bushes, strands) moved together in rotational or translational way. We assume that this is not the result of a coordinated movement, which would require sophisticated communication between filaments, but the result of the tight connection between filaments that are torn or pushed as a group. Filaments are apparently connected by an extracellular matrix based on a network of polysaccharide and proteins [39]. In our samples, a matrix could be observed by placing a needle tip into the medium next to filaments. A move of the needle caused a move of the nearby filaments in the same direction, although there was no direct contact between filament and needle. Through such a network, force could be generated between filaments for lateral movement, therefore there is no need for surface contact as in gliding motility.

For wild type and each mutant we made recordings of five or more different views and for each view about 5 recordings in series. Each recording was 30 s long. We tried to quantify movement speed or other parameters by different analysis software, but could not obtain reliable results. However, clear differences between strains became evident through qualitative comparisons. Here, we provide selected videos (movies 2-10) and merged images for each strain (Fig. 3). The merged images are composed of one picture at t=0 displayed in red and a second one taken after 10 s displayed in green. Moving filaments are thus red or green, all other positions of the image are grey.

**Figure 3.**
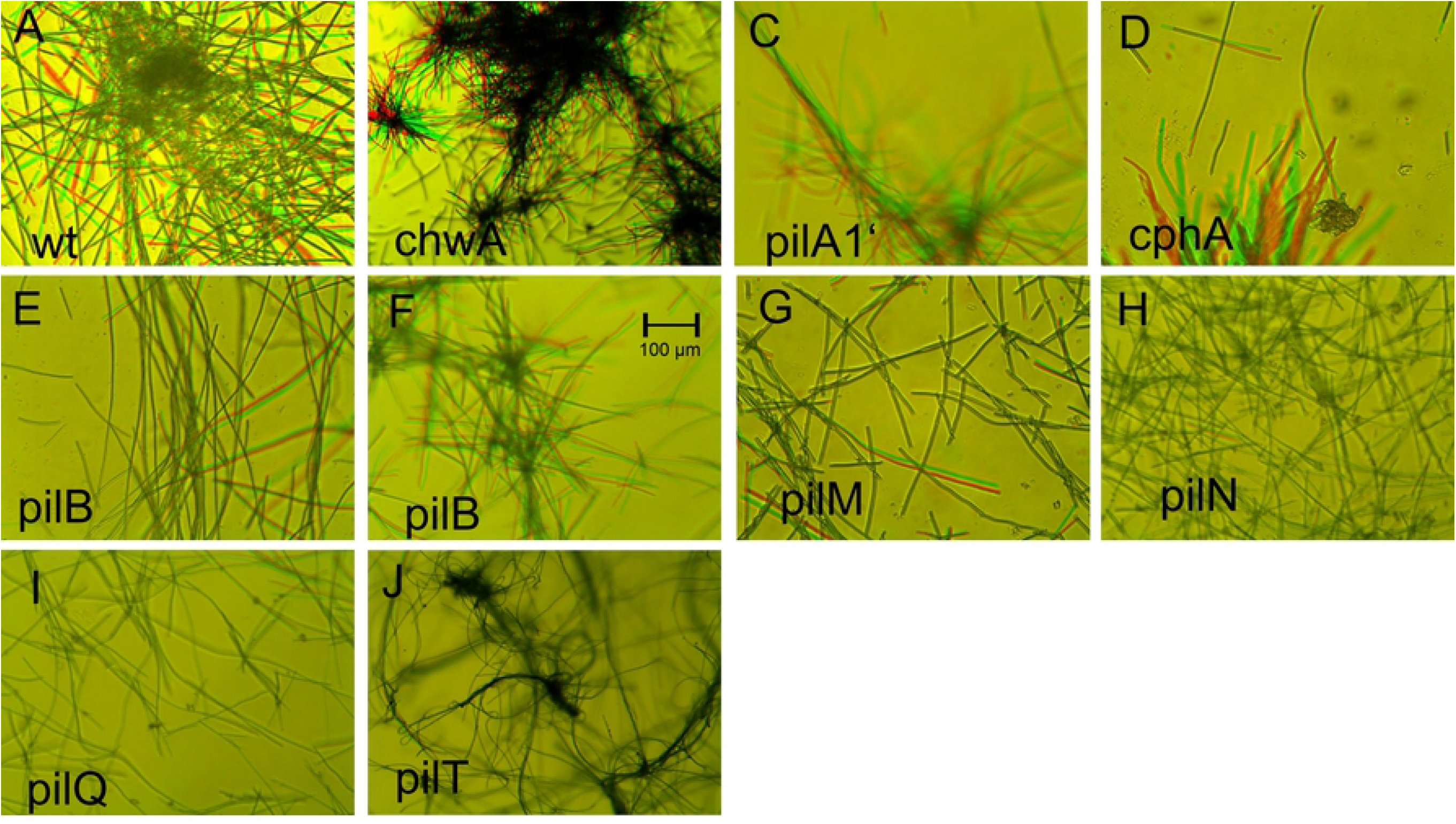
Motility in liquid culture. In each panel, two pictures with red and green false colors are superimposed. The time difference between both was 10 s. Moving filaments are highlighted by red or green color. The name of each mutant is given in the panels. For pilB there are two pictures, E and F, with weak and stronger movement, respectively.

Lateral movements of wild type filaments in liquid culture can be seen in movie 2. All filaments made lateral movements, although at a certain time point only a subfraction moved, as seen also in Fig. 3A. The *chwA* strain was indistinguishable from the wild type (Fig. 3B and movie 3). Also, *phrA’* and *cphA* displayed movements similar to the wild type (Fig. 3C and D, movies 4 and 5). An effect of the KanR cassette, as in the gliding motility experiments, was not obvious. The results of the *pilB* mutant differed qualitatively from one sample to another. In 3 of the 6 sample series, only few filaments moved with slow speed (as in Fig. 3 E). In the other 3 sample series, almost all filaments moved (as in Fig. 3 F and movie 6), although still less as compared to wild type. The reason for this difference among the *pilB* samples is not known. As in the gliding motility experiments, PilB is required for efficient movement, but dispensable for movement in general. With *pilM* mutants, we also observed that few filaments moved (Fig. 3 G), but most filaments remained silent. This result is again equivalent to the gliding motility study where very few filaments showed normal movement. For *pilN, pilQ* and *pilT* mutants, no lateral movements were observed. The *pil* mutants show clearly that type IV pili are required for lateral movement in liquid. The mechanism for lateral motility must be based on connections between filaments *via* the polysaccharide or protein extracellular network or from direct connections between filaments.

### Phototaxis

In the phototactic response, motile organisms move either towards the light or away from it. This response can most often be induced by unilateral light, *i.e.* by light that hits the organism within the area of possible moving directions. We performed many trials with *Phormidium lacuna* to induce phototactic movement by unilateral light (as outlined in Figs. 4 A, B) but the filaments remained equally distributed in the Petri dish (as in Fig. 4 E). Neither the use of agar plates nor the use of different irradiation wavelengths nor the variation of intensities resulted in a directional response. However, in time lapse studies with *Phormidium filaments* we observed that filaments sometimes gather in the white microscope light that hits the sample from below. An example for this effect is shown in movie 1. We therefore suggested that light projected vertically to the movement area can induce phototaxis of *Phormidium lacuna*. Directional effects by vertical light have been described for other organisms such as the alga *Euglena gracilis* [40] or *Phormidium uncinatum* [41, 42]. In our subsequent studies, *Phormidium lacuna* filaments were irradiated with light emitting diodes (LEDs) from below. In standard assays, 5 cm Petri dishes (without agar) that were filled with 8 ml Utraturrax-treated filaments in f/2 medium were placed on a holder in which a light emitting diode shines light through a short tunnel on the specimen from below, at the position of the center of the Petri dish (Figs 4C, D). We observed that red light, blue light and green light of around 5 −50 μmol m^−2^ s^−1^ induce a clear gathering of wild type filaments towards the light in the center (Figs 4 F, G, H). The effect could be observed after several hours but increased during 1 or 2 d. Thus, the movement of *Phormidium lacuna* in light follows rather an intensity gradient than the light direction and behaves different from single celled cyanobacteria. Responses within light gradients have been termed phototaxis [40], like the directional movement towards unilateral light, or, more specifically, phobotaxis [42]. We prefer to use the more universal term phototaxis here, because this term expresses better that filaments move to the light. It should be stressed that this movement takes place in Petri dishes with or without agar and that the majority of phototaxis experiments were performed without agar.

**Figure 4.**
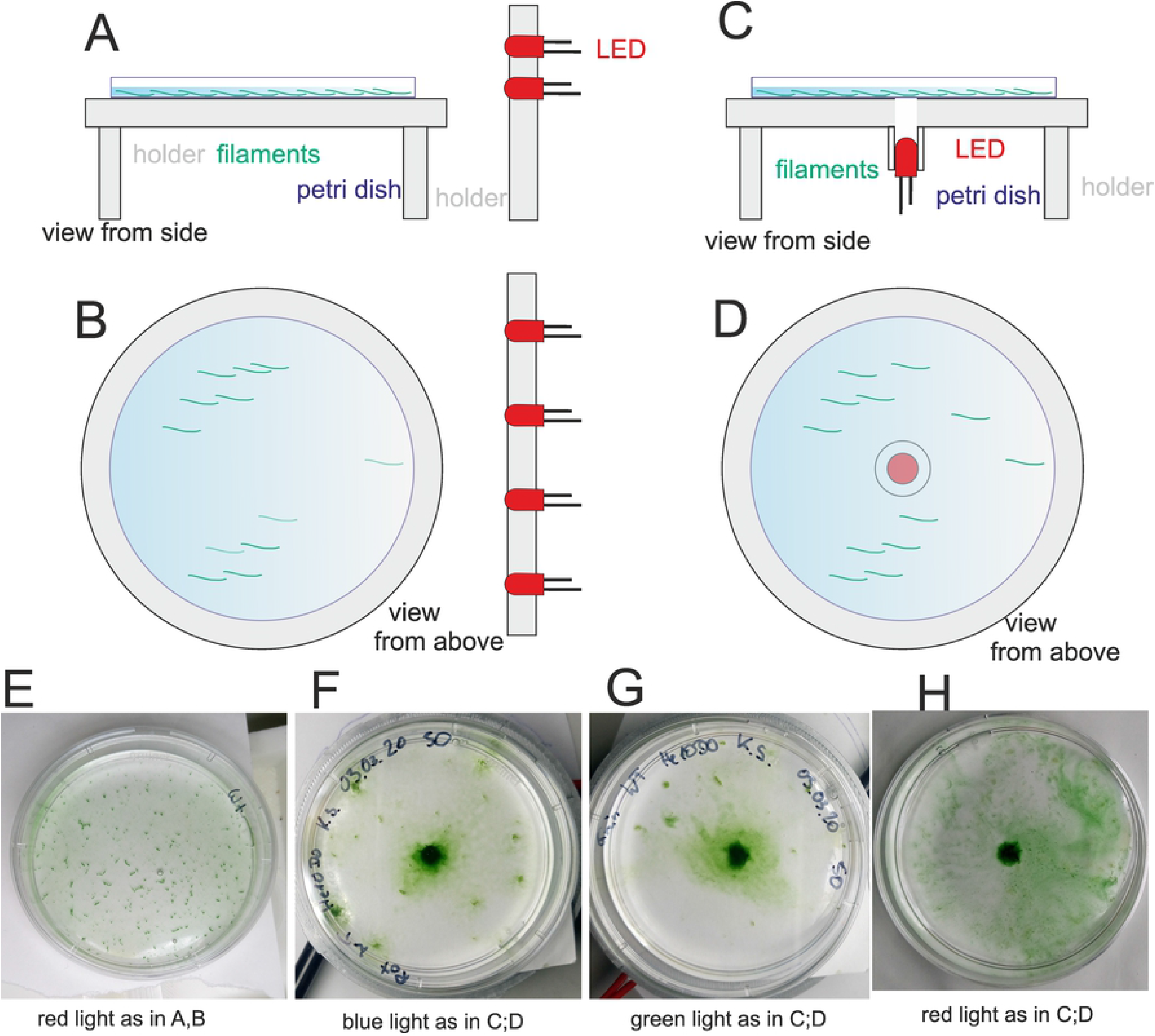
Setup of phototaxis experiments with unilateral irradiation (from the side, A, B) or vertical irradiation (C,D) and examples for irradiations with red light from the side (E) or vertical blue (F), green (G) or red light (H). Light intensities were 25 μmol m^−2^ s^−1^ for blue (450 nm) and green (550 nm) and 15 μmol m^−2^ s^−1^ for red (655 nm). The duration of irradiation was 2 d.

Comparative analyses of wild type and mutants were performed with red light at a light intensity of 15 μmol m^−2^ s^−1^ for 2 d. Thereafter, a photograph was recorded from the Petri dish. We made routinely made a second photograph after a gentle shaking of the Petri dish. The diameter of the covered circle was measured between the two positions of half maximal pixel intensity along a line through the center of the circle (Fig. 5 A, B), this value was used to calculate the circle area. The area value stands as a measure for the magnitude of the phototactic response. Shaking reduced the size of the densely covered area. In this way, the data distinguish between all filaments that have moved to the center and those filaments that were more tightly bound to the surface. Shaking did not increase the error values to a relevant extent (Fig. 5 C), which supports the reliability of the approach.

**Figure 5.**
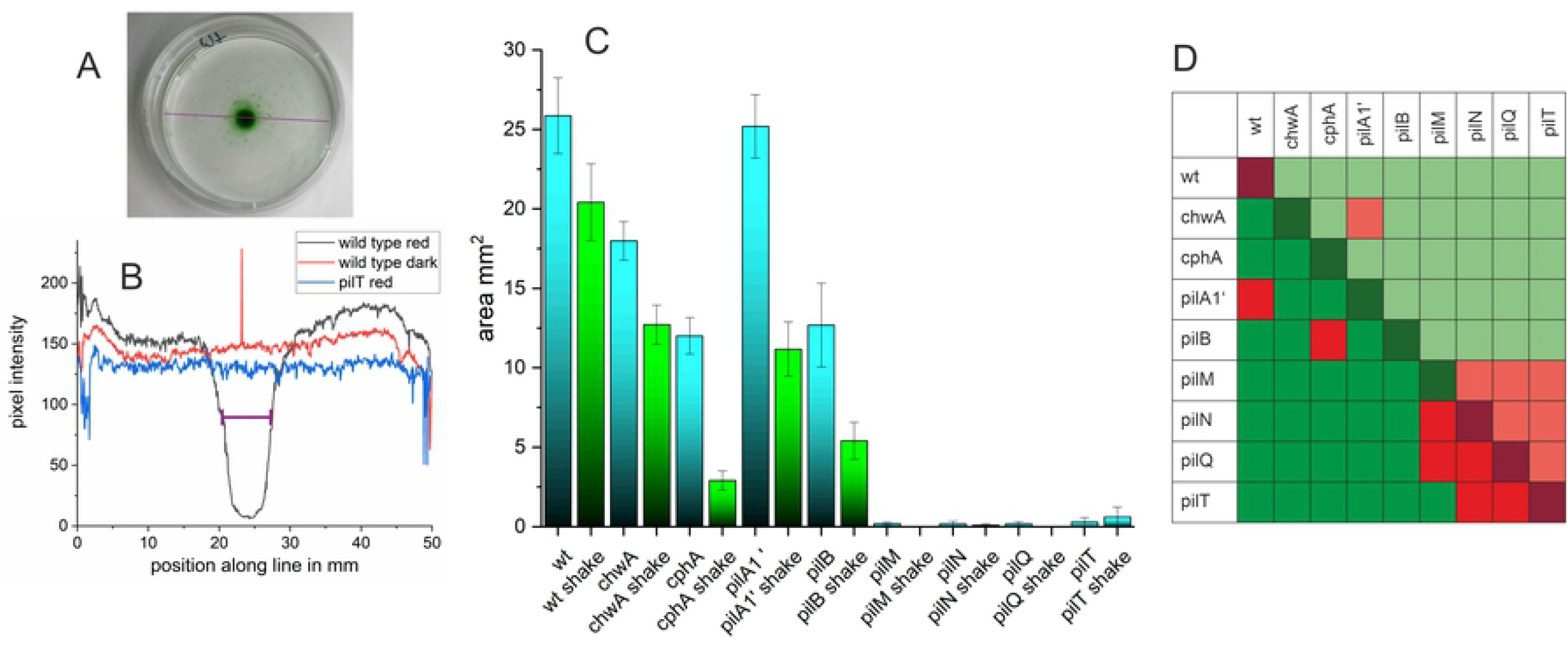
Phototaxis experiments with *Phormidium lacuna* wild type and various mutants. (A) Petri dish with wild type filaments after 2 d irradiation with central LED from below. Pixel intensities along the violet line are taken for calculation of the diameter. (B) Typical profiles of pixel intensities as outlined in A for wild type after illumination, wild type without illumination and pilT after illumination. The estimated diameter for wild type red is given by violet distance (C) Mean area after 2 d in 5 mm LED, before (blue bars) and after (green bars) shaking. (D) Kruskal Wallis significance test, comparison of all samples with each other. The lower left triangle shows samples before shaking, the upper right triangle shows comparisons of samples after shaking, and the diagonal shows the comparisons of each samples before and after shaking. Green squares indicate significance with p < 0.05, red squares indicate non significant differences.

The circle area of the light beam is 25 mm^2^ and in a clear response the entire area is completely covered with filaments. We experienced that a response could result in a covered area even larger than 25 mm^2^, and any smaller covered areas were also often observed. In samples where no accumulation in the center is obvious, as in the dark control or *pilT*^-^ examples shown in Fig. 5 B, we set the area value to 0. The data processing showed a broad distribution of covered areas (supplemental Figure S1) for wild type, *cphA, pilA1’, chwA, pilB*. The ranges were from no response to intermediate and strong responses with covered areas above 25 mm^2^. The mean values are presented in Fig. 5 C. We estimated the significances of differences between all possible pairs of samples. Since the values of the covered areas are not normally distributed, significances were estimated according to Kruskal and Wallis [37]. Significance of sample pairs before shaking is presented by colored squares in the lower left triangle of Fig. 5D, the pairs of samples after shaking are presented by the colored squares in the upper right triangle of Fig. 5D and the significance of each sample before and after shaking are given by the colored squares in the diagonal in of Fig. 5D.

The wild type covered an average area of 25 mm^2^. Shaking reduced the wild type area to a mean value of 20 mm^2^, but the difference before and after shaking was not significant. With ca. 18 mm^2^, the mean covered area of *chwA* was reduced as compared to the wild type; shaking reduced the area to 12 mm^2^, this reduction was significant. Like in the motility-on-agar experiments we assume that the reduced response of *chwA* vs. wild type results from the KanR cassette.

In case of *pilA1‘*, the covered area was 25 mm^2^, just as large as that of the wild type. Since *pilA1’* also has a KanR cassette, we assume that the partial loss of *pilA1* gene copies results in a positive effect on phototaxis by which the negative effect of KanR is compensated. Note that it is to be expected that *pilA1’* produces less PilA1 filaments than the wild type. After shaking, *pilA1’* covered only 12 mm^2^, comparable to *chwA*.

The response of *cphA* was further reduced compared to wild type, *chwA* and *pilA1’*. Before and after shaking the covered areas were 12 mm^2^ and 2 mm^2^, respectively. These low values of *cphA* imply that in the wild type, CphA regulates the phototaxis response as a photoreceptor.

The *pilB* mutant had a covered area of 12 mm^2^ and an area of 5 mm^2^ after shaking. The pattern is comparable with *cphA*. The cause for reduction could be the reduced motility of *pilB*, but also a reduced adhesion, which could result from partially defective type IV pili.

The other pilin mutants *pilN, pilM, pilQ and pilT* had no detectable phototactic response. Only in exceptional cases, a response was obtained. That for a phototactic response the integrity of type IV pili is required is no surprise, as phototaxis without motion is not possible.

## Discussion

We describe here for the first time mutant effects on the motility of a member of the Oscillatoriales, *Phormidium lacuna*. The motility of the genera *Oscillatoria* or *Phormidium* have been scientifically investigated since the 19^th^ century [6, 41, 43–45]. Cellular and molecular studies on motility of cyanobacteria have, however, concentrated on the model organism *Synechocystis* PCC 6803 [9, 46], a unicellular species. Few experiments followed with *Nostoc punctiforme* (Nostocales) and *Synechococcus elongatus* (single celled) [11]. There are three major differences between motilities of *Phormidium lacuna* and single celled cyanobacteria. (i) The gliding motility of *Phormidium lacuna* is characterized by continued change of direction, combined with bending movements, and thus essentially different from the straight movement of single celled cyanobacteria. (ii) The lateral motion of *Phormidium lacuna* that has been observed in liquid medium (movies 2-10) cannot work in single celled species. (iii) Finally, the phototactic movement of *Phormidium lacuna* is induced by light intensity gradients, in contrast to the movement towards unilateral light of single celled species. The mechanisms of light perception and direction finding must be completely different between these species. Although type IV pili are widely distributed in cyanobacteria and present in *Phormidium lacuna*, it was initially not clear whether these pili are involved in motions of *Phormidium lacuna*. Lateral motion, independent of surface contacts, is rarely described in cyanobacteria. The direction of lateral movement is opposed to the longitudinal direction in gliding motility. Because *pilM, pilN, pilQ* and *pilT* were neither motile in gliding motility nor in lateral movements, both kinds of movement must be driven by type IV pili. For lateral motility, this implies that the outer ends of the pili are connected with the extracellular matrix or with pili of other cells, in order to generate force for the motions. What evolutionary advantage can be found behind lateral motions in liquid? The back and forth movements do not allow the filaments to reach another place. However, filaments join neighboring filaments and form aggregates, and this kind of biofilm formation could protect against predators. The counterpart of these aggregates are the aligned filaments on agar that form leaflike structures (Fig. 1). Also in this case, the getting together could result in protection against predators. Type IV pili play also an important role in the natural transformation process [22]. DNA from the medium is bound to PilA and torn into the intramembrane space, converted to single stranded DNA and brought by other proteins into the interior of the cell [47]. Since the PilA pili are always expelled and rejected, PilA bound DNA could as well be transported outward. DNA fragments that are *e.g*. formed during replication could bind to PilA at the plasma membrane. In this way, continuous exchange of DNA could be possible between filaments that are connected by type IV pili and a close proximity between filaments would increase the efficiency.

The results from all three assays, analyzing gliding motility on agar, lateral motions in liquid and phototaxis indicate that PilM, PilN, PilQ and PilT are essential for the function of the type IV pili – without these any kind of motion is inhibited. PilN and PilM are pore forming components in the inner membrane and PilQ is involved in pore forming at the outer membrane. PilT is an ATPase for retraction of the pilus. Clearly, the type IV pilus is dysfunctional if any single compound is missing. We would like to stress here that it is very unlikely that the immotile phenotype results from second site mutations. We have meanwhile performed many *Phormidium* transformations at other sites. The immobile phenotype was never observed.

In *pilB* mutants motility is only partially blocked in the three motion assays. PilB is the ATPase for pilus expelling. The PCR results indicate complete segregation of the interrupted gene. As the other mutants, this line was kept continuously on Kan medium until it was used for experiments. There is thus no reason to assume that wild type *pilB* copies are still abundant in the cells. We also checked whether PilB homologs are present in *Phormidium lacuna*, but found no indication for such redundancy. We therefore must assume that the expelling of the pilus is driven by an alternative mechanism. One possible explanation is that PilT serves as ATPase for inward and outward directions.

For *pilA1* we did only obtain partially segregated disruption lines, and in case of *pilD* it was even not possible to select lines in which the disrupted gene is partially segregated. The loss of PilD in *Synechocystis* PCC 6803 is lethal because PilA1 prepilin proteins accumulate in the plasma membrane and induce toxic effects [36]. A *pilA1* mutant would, however, not cause such effects. It is difficult to understand why *Phormidium lacuna* cells cannot survive without *pilA1. Phormidium lacuna* has 2 homologs of PilA2 (WP_140409148, 30% identities of overlapping region, WP_087711780.1, 31%), but no other PilA homologs. For *Phormidium lacuna* we could imagine that misassembly of the pilus due to lack of PilA1 results in accumulation of PilA2 proteins in the plasma membrane and in cell death of completely segregated *pilA1* mutants.

The *cphA* mutant shows motility on agar comparable with the wild type, *chwA* and *pila1*. The phototaxis response of *cphA* was, however, clearly diminished compared to the other three lines. The *cphA* effect is thus specific for phototaxis. We therefore assume that CphA operates as a phototaxis photoreceptor in *Phormidium lacuna*. Typical phytochromes are red light sensors and our relevant phototaxis experiments have been performed in red light. There is, however, a weak phototaxis response in the *cphA* mutant. This must be regulated by a different photoreceptor such as a cyanobacteriochrome like PixJ or a photoreceptor with a BLUF domain. Phototaxis of *Phormidium lacuna* is observed in blue, green and red light. Therefore it is very likely that more than one single photoreceptor modulates phototaxis. The role of CphA in phototaxis is another major difference between *Synechocystis* PCC 6803 and *Phormidium lacuna*, since knockout mutants of the homologous Cph1 in *Synechocystis* were not affected in phototaxis. Although cyanobacterial phytochrome Cph1was discovered as one of the first cyanobacterial photoreceptors [48, 49], the biological role of cyanobacterial phytochromes remained obscure.

We assume that phototaxis of *Phormidium lacuna* is a result of random movement as e.g. seen in movie 1 and light-dependent attachment to the surface. It is also possible that the filaments follow a light gradient that is formed by stray light from the vertically projecting LEDs. However, the lack of phototaxis in unilateral light speaks against this possibility. How does a filament decide to switch from mobile stage to attachment? The surface attachment in the area of the light beam is only found if the light irradiates a certain region of the Petri dish but not if the entire container is irradiated, e.g. in the growth chamber. Therefore, the attachment to the surface must be triggered by temporal or spatial “step up” of light intensity. For spatial gradient detection, comparison of light intensities at both apical ends of the filament is required. In the center of the LED light beam there is no light gradient, but filaments attach here too. Moreover, filaments must recognize neighboring filaments in order to obtain optimal horizontal distribution. Intracellular communication and macroscopic pattern recognition show that the phototaxis system of *Phormidium lacuna* is more complex than that of *Synechocystis* PCC6803 or other single celled cyanobacteria.

## Author Contributions

Generation of mutants: NoW, KF, MM, FZ, JB, NS; VA; motility experiments: JB, KF, MM, NaW, KS; data curation, manuscript writing, coordination: TL.

## Supplementary Material

Supplementary Table 1, used primers

Supplementary Figure 1, PCR results

Supplemental Figure 2, histogram of phototaxis

Movie 1 Movement of *Phormidium lacuna* wild type filaments after Ultraturrax treatment on agar, time lapse with 1 image every min; 10 x objective.

Movie 2 Movement of *Phormidium lacuna* wild type filaments in liquid medium after Ultraturrax treatment; 10 x objective.

Movie 3 Movement of *Phormidium lacuna chwA* filaments in liquid medium after Ultraturrax treatment; 10 x objective.

Movie 4 Movement of *Phormidium lacuna cphA* filaments in liquid medium after Ultraturrax treatment; 10 x objective.

Movie 5 Movement of *Phormidium lacuna pilA1’* filaments in liquid medium after Ultraturrax treatment; 10 x objective.

Movie 6 Movement of *Phormidium lacuna pilB* filaments in liquid medium after Ultraturrax treatment; 10 x objective.

Movie 7 Movement of *Phormidium lacuna pilM* filaments in liquid medium after Ultraturrax treatment; 10 x objective.

Movie 8 Movement of *Phormidium lacuna pilN* filaments in liquid medium after Ultraturrax treatment; 10 x objective.

Movie 9 Movement of *Phormidium lacuna pilQ* filaments in liquid medium after Ultraturrax treatment; 10 x objective.

Movie 10 Movement of *Phormidium lacuna pilT* filaments in liquid medium after Ultraturrax treatment; 10 x objective.

## References

1. Schirrmeister BE, Gugger M, Donoghue PCJ. Cyanobacteria and the Great Oxidation Event: evidence from genes and fossils. Palaeontology. 2015;58(5):769–85. doi: 10.1111/pala.12178. PubMed PMID: WOS:000360586100002.

2. Margulis L. The origin of plant and animal cells. Am Sci. 1971;59(2):230–5.

3. Delwiche CF, Palmer JD. The origin of plastids and their spread via secondary symbiosis. Plant Systematics and Evolution. 1997:53–86. PubMed PMID: WOS:000071339300004.

4. Castle ES. Observations on motility in certain Cyanophyceae. Biol Bull Marine Biol Lab. 1926;51((2)):69–72. doi: 10.2307/1536537. PubMed PMID: BIOSIS:PREV19270100003415.

5. Lazaroff N, Schiff J. Action spectum for developmental photoinduction of the blue-green alga *Nostoc muscorum*. Science. 1962;137:603–4.

6. Nultsch W. [Effect of redox systems on the motility and the phototactic reactions of Phormidium uncinatum]. Arch Mikrobiol. 1968;63(4):295–320. Epub 1968/01/01. PubMed PMID: 5709367.

7. Komarek J, Kastovsky J, Mares J, Johansen JR. Taxonomic classification of cyanoprokaryotes (cyanobacterial genera) 2014, using a polyphasic approach. Preslia. 2014;86(4):295–335. PubMed PMID: WOS:000348630000001.

8. Khayatan B, Meeks JC, Risser DD. Evidence that a modified type IV pilus-like system powers gliding motility and polysaccharide secretion in filamentous cyanobacteria. Molecular Microbiology. 2015;98(6):1021–36.

9. Bhaya D, Bianco NR, Bryant D, Grossman A. Type IV pilus biogenesis and motility in the cyanobacterium Synechocystis sp. PCC6803. Mol Microbiol. 2000;37:941–51.

10. Fiedler B, Borner T, Wilde A. Phototaxis in the cyanobacterium Synechocystis sp. PCC 6803: role of different photoreceptors. Photochem Photobiol. 2005;81(6):1481–8. Epub 2005/12/16. doi: 10.1562/2005-06-28-RA-592. PubMed PMID: 16354116.

11. Yang Y, Lam V, Adomako M, Simkovsky R, Jakob A, Rockwell NC, et al. Phototaxis in a wild isolate of the cyanobacterium Synechococcus elongatus. Proc Natl Acad Sci U S A. 2018;115(52):E12378–E87. Epub 2018/12/16. doi: 10.1073/pnas.1812871115. PubMed PMID: 30552139; PubMed Central PMCID: PMCPMC6310787.

12. Wall D, Kaiser D. Type IV pili and cell motility. Molecular Microbiology. 1999;32(1):1–10. doi: 10.1046/j.1365-2958.1999.01339.x. PubMed PMID: WOS:000079767100001.

13. Souza ID, Maissa N, Ziveri J, Morand PC, Coureuil M, Nassif X, et al. Meningococcal disease: A paradigm of type-IV pilus dependent pathogenesis. Cellular Microbiology. 2020;22(4). doi: 10.1111/cmi.13185. PubMed PMID: WOS:000521198000011.

14. Nudleman E, Kaiser D. Pulling Together with Type IV Pili. Journal of Molecular Microbiology and Biotechnology. 2004;7(1-2):52–62. doi: 10.1159/000077869.

15. Carbonnelle E, Helaine S, Nassif X, Pelicic V. A systematic genetic analysis in Neisseria meningitidis defines the Pil proteins required for assembly, functionality, stabilization and export of type IV pili. Molecular Microbiology. 2006;61(6):1510–22. doi: 10.1111/j.1365-2958.2006.05341.x. PubMed PMID: WOS:000240368000014.

16. Berry JL, Pelicic V. Exceptionally widespread nanomachines composed of type IV pilins: the prokaryotic Swiss Army knives. Fems Microbiology Reviews. 2015;39(1):134–54. doi: 10.1093/femsre/fuu001. PubMed PMID: WOS:000351353000008.

17. Cengic I, Uhlen M, Hudson EP. Surface Display of Small Affinity Proteins on Synechocystis sp Strain PCC 6803 Mediated by Fusion to the Major Type IV Pilin PilA1. Journal of Bacteriology. 2018;200(16):19. doi: 10.1128/jb.00270-18. PubMed PMID: WOS:000439777600014.

18. Conradi FD, Zhou RQ, Oeser S, Schuergers N, Wilde A, Mullineaux CW. Factors Controlling Floc Formation and Structure in the Cyanobacterium Synechocystis sp. Strain PCC 6803. J Bacteriol. 2019;201(19). Epub 2019/07/03. doi: 10.1128/JB.00344-19. PubMed PMID: 31262837; PubMed Central PMCID: PMCPMC6755745.

19. Taton A, Erikson C, Yang Y, Rubin BE, Rifkin SA, Golden JW, et al. The circadian clock and darkness control natural competence in cyanobacteria. Nature Communications. 2020;11(1). doi: 10.1038/s41467-020-15384-9.

20. Yoshihara S, Geng X, Okamoto S, Yura K, Murata T, Go M, et al. Mutational analysis of genes involved in pilus structure, motility and transformation competency in the unicellular motile cyanobacterium Synechocystis sp. PCC 6803. Plant Cell Physiol. 2001;42(1):63–73. Epub 2001/02/07. doi: 10.1093/pcp/pce007. PubMed PMID: 11158445.

21. Wendt KE, Pakrasi HB. Genomics Approaches to Deciphering Natural Transformation in Cyanobacteria. Frontiers in Microbiology. 2019;10:7. doi: 10.3389/fmicb.2019.01259. PubMed PMID: WOS:000470976200002.

22. Nies F, Mielke M, Pochert J, Lamparter T. Natural transformation of the filamentous cyanobacterium Phormidium lacuna. PLoS One. 2020;15(6):e0234440. Epub 2020/06/13. doi: 10.1371/journal.pone.0234440. PubMed PMID: 32530971; PubMed Central PMCID: PMCPMC7292380.

23. Read N, Connell S, Adams DG. Nanoscale visualization of a fibrillar array in the cell wall of filamentous cyanobacteria and its implications for gliding motility. J Bacteriol. 2007;189(20):7361–6. Epub 2007/08/19. doi: 10.1128/JB.00706-07. PubMed PMID: 17693519; PubMed Central PMCID: PMCPMC2168455.

24. Wilde A, Fiedler B, Borner T. The cyanobacterial phytochrome Cph2 inhibits phototaxis towards blue light. Mol Microbiol. 2002;44(4):981–8. Epub 2002/05/16. doi: 10.1046/j.1365-2958.2002.02923.x. PubMed PMID: 12010493.

25. Schuergers N, Nurnberg DJ, Wallner T, Mullineaux CW, Wilde A. PilB localization correlates with the direction of twitching motility in the cyanobacterium Synechocystis sp. PCC 6803. Microbiology (Reading). 2015;161(Pt 5):960–6. Epub 2015/02/28. doi: 10.1099/mic.0.000064. PubMed PMID: 25721851.

26. Schuergers N, Lenn T, Kampmann R, Meissner MV, Esteves T, Temerinac-Ott M, et al. Cyanobacteria use micro-optics to sense light direction. Elife. 2016;5. Epub 2016/02/10. doi: 10.7554/eLife.12620. PubMed PMID: 26858197; PubMed Central PMCID: PMCPMC4758948.

27. Lamparter T, Esteban B, Hughes J. Phytochrome Cph1 from the cyanobacterium *Synechocystis* PCC6803: purification, assembly, and quaternary structure. Eur J Biochem. 2001;268:4720–30.

28. Ikeuchi M, Ishizuka T. Cyanobacteriochromes: a new superfamily of tetrapyrrole-binding photoreceptors in cyanobacteria. Photochem Photobiol Sci. 2008;7(10):1159–67.

29. Schirrmeister BE, Antonelli A, Bagheri HC. The origin of multicellularity in cyanobacteria. Bmc Evolutionary Biology. 2011;11. doi: 10.1186/1471-2148-11-45. PubMed PMID: WOS:000289411400001.

30. Nies F, Worner S, Wunsch N, Armant O, Sharma V, Hesselschwerdt A, et al. Characterization of Phormidium lacuna strains from the North Sea and the Mediterranean Sea for biotechnological applications. Process Biochemistry. 2017;59:194–206. doi: 10.1016/j.procbio.2017.05.015. PubMed PMID: WOS:000412376700009.

31. Guillard RR, Ryther JH. Studies of marine planktonic diatoms. I. Cyclotella nana Hustedt, and Detonula confervacea (cleve) Gran. Can J Microbiol. 1962;8:229–39.

32. Hübschmann T, Yamamoto H, Gieler T, Murata N, Börner T. Red and far-red light alter the transcript profile in the cyanobacterium Synechocystis sp. PCC 6803: impact of cyanobacterial phytochromes. FEBS Lett. 2005;579(7):1613–8.

33. Herdman M, Coursin T, Rippka R, Houmard J, Tandeau de Marsac N. A new appraisal of the prokaryotic origin of eukaryotic phytochromes. J Mol Evol. 2000;51:205–13.

34. Yoshihara S, Ikeuchi M. Phototactic motility in the unicellular cyanobacterium Synechocystis sp. PCC 6803. Photochem Photobiol Sci. 2004;3(6):512–8.

35. Nölling J, Breton G, Omelchenko MV, Makarova KS, Zeng Q, Gibson R, et al. Genome sequence and comparative analysis of the solvent-producing bacterium Clostridium acetobutylicum. J Bacteriol. 2001;183(16):4823–38. Epub 2001/07/24. doi: 10.1128/jb.183.16.4823-4838.2001. PubMed PMID: 11466286; PubMed Central PMCID: PMCPMC99537.

36. Linhartova M, Bucinska L, Halada P, Jecmen T, Setlik J, Komenda J, et al. Accumulation of the Type IV prepilin triggers degradation of SecY and YidC and inhibits synthesis of Photosystem II proteins in the cyanobacterium Synechocystis PCC 6803. Molecular Microbiology. 2014;93(6):1207–23. doi: 10.1111/mmi.12730. PubMed PMID: WOS:000342757200011.

37. Kruskal WH, Wallis WA. Use of Ranks in One-Criterion Variance Analysis. Journal of the American Statistical Association. 1952;47(260):583–621. doi: 10.1080/01621459.1952.10483441.

38. Crozier WJ, Federighi H. CRITICAL THERMAL INCREMENT FOR THE MOVEMENT OF OSCILLATORIA. The Journal of general physiology. 1924;7(1):137–50. doi: 10.1085/jgp.7.1.137. PubMed PMID: MEDLINE:19872119.

39. Ge HM, Xia L, Zhou XP, Zhang DL, Hu CX. Effects of Light Intensity on Components and Topographical Structures of Extracellular Polysaccharides from the Cyanobacteria Nostoc sp. J Microbiol. 2014;52(2):179–83. doi: 10.1007/s12275-014-2720-5. PubMed PMID: WOS:000330416200012.

40. Diehn B. PHOTOTAXIS AND SENSORY TRANSDUCTION IN EUGLENA. Science. 1973;181(4104):1009–15. doi: 10.1126/science.181.4104.1009. PubMed PMID: WOS:A1973Q569300011.

41. Nultsch W, Hader D. Bestimmung der phototaktischen Unterschiedsschwelle bei *Phormidium uncinatum*. Ber Dt Bot Ges 83,185-192. 1970; 83(5/6):185–92.

42. Hader DP. PARTICIPATION OF 2 PHOTOSYSTEMS IN PHOTO-PHOBOTAXIS OF PHORMIDIUM-UNCINATUM. Archives of Microbiology. 1974;96(3):255–66. doi: 10.1007/bf00590181. PubMed PMID: WOS:A1974S475900008.

43. Vincent BdS. Oscillaire. Dictionnaire Classique d’Histoire Naturelle Tome 121827.

44. Burkholder PR. Movement in the Cyanophyceae. The Quarterly Review of Biology. 1934;9(4):438–59.

45. Nultsch W. [On the antagonism of atabrine and flavine nucleotides in the behavior of movement and reaction to light of Phormidium uncinatum]. Arch Mikrobiol. 1966;55(2):187–99. Epub 1966/11/11. PubMed PMID: 5992184.

46. Bhaya D, Takahashi A, Grossman AR. Light regulation of type IV pilus-dependent motility by chemosensor-like elements in Synechocystis PCC6803. Proc Natl Acad Sci U S A. 2001;98(13):7540–5.

47. Chen I, Dubnau D. DNA uptake during bacterial transformation. Nature Reviews Microbiology. 2004;2(3):241–9. doi: 10.1038/nrmicro844. PubMed PMID: WOS:000220431800015.

48. Hughes J, Lamparter T, Mittmann F, Hartmann E, Gärtner W, Wilde A, et al. A prokaryotic phytochrome. Nature. 1997;386:663-.

49. Yeh KC, Wu SH, Murphy JT, Lagarias JC. A cyanobacterial phytochrome two-component light sensory system. Science. 1997;277(5331):1505–8.

50. Needleman SB, Wunsch CD. A general method applicable to the search for similarities in the amino acid sequence of two proteins. Journal of Molecular Biology. 1970;48(3):443–53. doi: https://doi.org/10.1016/0022-2836(70)90057-4.

